## Extended Data including Supplementary Figures for "Pathway choice in the alternative telomere lengthening in neoplasia is dictated by replication fork processing mediated by EXD2’s nuclease activity"

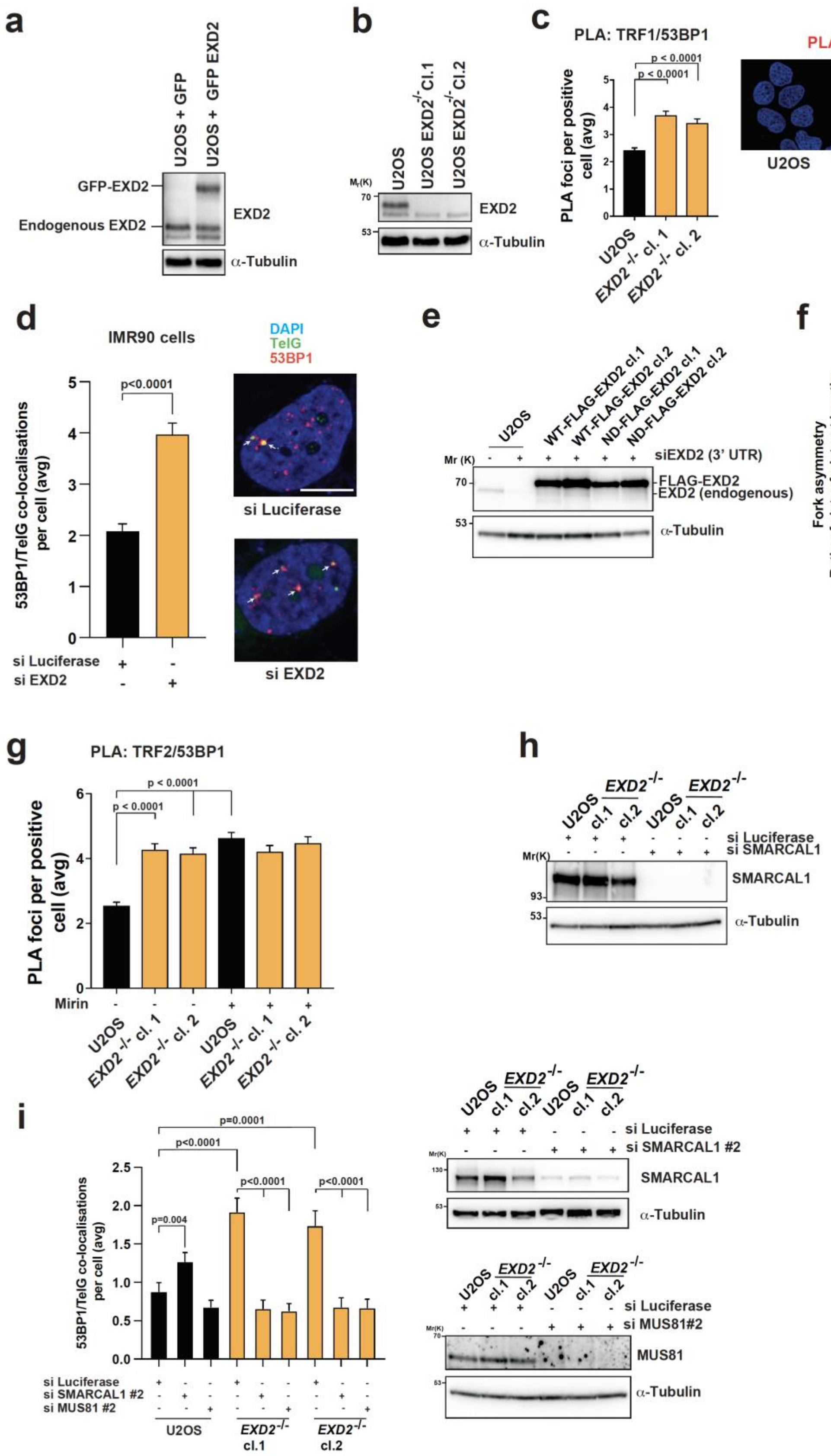


**Supplementary Figure 1**

**Supplementary Figure 1**

a) Western blot confirming expression of GFP-EXD2, endogenous and ectopically expressed GFP-EXD2 are indicated, $\alpha$-Tubulin acts as a loading control.

b) Western blot confirming EXD2 knockout in U2OS cells, $\alpha$-Tubulin acts as a loading control.

c) Quantification of PLA foci per focus positive nucleus using antibodies recognising TRF1 (a shelterin component) and 53BP1 (a DSB marker) in WT U2OS or *EXD2*$^{-/-}$ cells (n= at least 220 cells from 3 independent experiments, statistical significance was determined by Mann-Witney analysis, bars represent +/-SEM, scale bar = 10$\mu$m).

d) Immuno-FISH staining in IMR90 cells treated with control siRNA (si Luciferase) or siRNA targeting EXD2, as indicated. 53BP1 acts as a DSB marker, TelG PNA FISH probe staining acts as a marker for the telomere and DAPI acts as a nuclear stain (n= at least 161 cells from 3 independent experiments, statistical analysis by Mann-Witney test, bars represent +/- SEM, scale bar = 10$\mu$m).

e) Confirmation of endogenous EXD2 knockdown in U2OS WT cells and cells recomplemented with either WT or nuclease dead (ND) mutant FLAG- EXD2. $\alpha$-Tubulin acts as a loading control.

f) DNA fiber analysis of global replication fork asymmetry in WT U2OS or *EXD2*$^{-/-}$ cells. CldU/IdU labelling strategy is indicated in the schematic (n= 3 independent experiments, statistical significance was determined by Mann-Witney test, boxes represent 5-95 percentile).

g) Quantification of PLA foci per focus positive nucleus using antibodies recognising TRF2 and 53BP1 in WT U2OS or *EXD2*$^{-/-}$ cells treated with Mirin or DMSO, as indicated (n= at least 230 cells from 3 independent experiments, statistical significance was determined by Mann-Witney test, bars represent +/- SEM).

h) Western blotting confirming SMARCAL1 knockdown in WT U2OS or *EXD2*$^{-/-}$ cells treated with either control siRNA or siRNA targeting SMARCAL1, as indicated. $\alpha$-Tubulin acts as a loading control.

i) Quantification of 53BP1/TelG co-localisations by immuno-FISH staining of cells treated with control siRNA (si Luciferase) or siRNA targeting MUS81 or SMARCAL1, as indicated (n= 100 cells from 2 independent experiments, statistical analysis by Mann-Witney test). Confirmation of knockdown of SMARCAL1 and MUS81 by western blotting is presented. $\alpha$-Tubulin acts as a loading control.

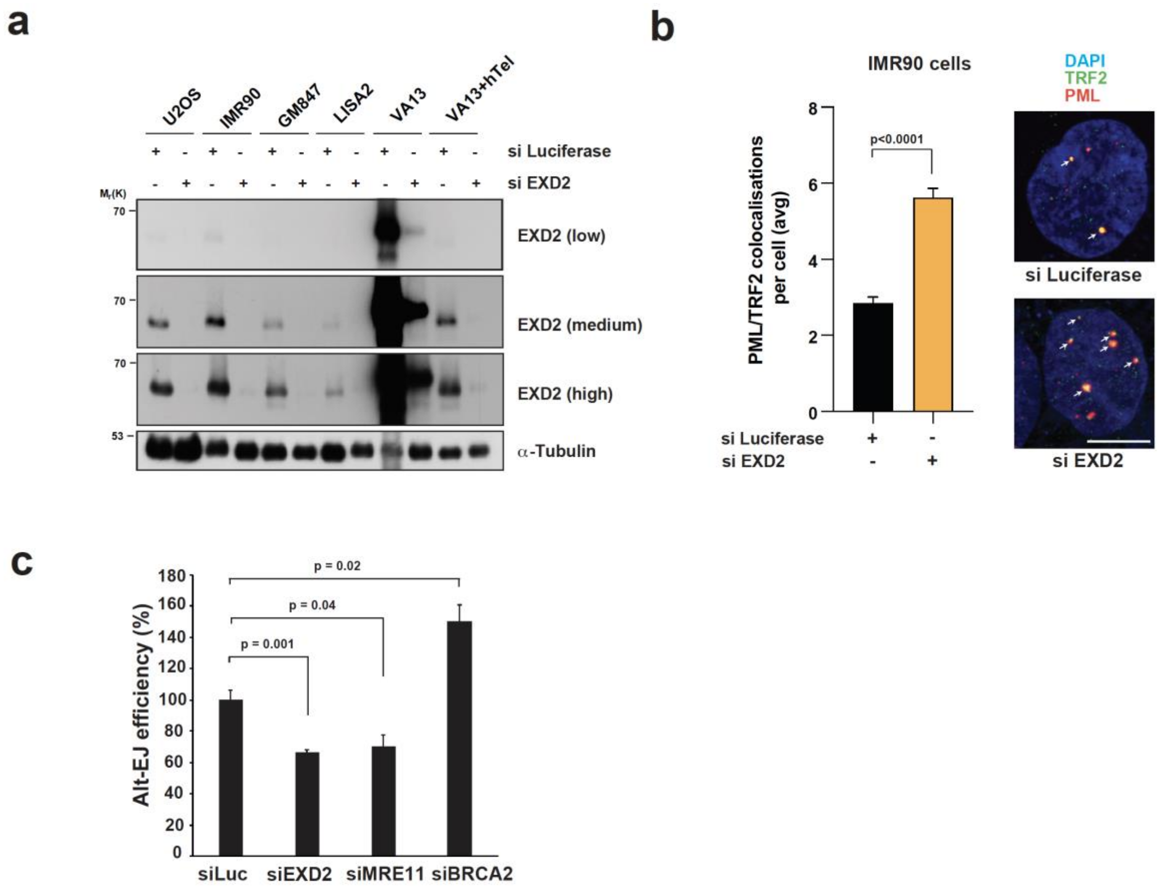


Supplementary Figure 2

**Supplementary Figure 2**

a) Confirmation of EXD2 knockdown in a panel of ALT+ cells (VA-13+hTel acts as a negative control); multiple exposures are presented (low, medium, high).α-Tubulin acts as a loading control.

b) Quantification of the incidence of ALT-associated PML bodies in IMR90 cells treated with either control siRNA (si Luciferase) or si RNA targeting EXD2, as indicated by immunofluorescence staining using antibodies against TRF2 (red) and PML (green), DAPI acts as a nuclear stain (n= at least 199 cells from 3 independent experiments, statistical significance confirmed by Mann-Witney test, bars represent +/- SEM, scale bar = 10μm).

c) ALT-EJ reporter assay carried out in U2OS cells treated with control siRNA or siRNA targeting EXD2 or BRCA2 (positive control for this assay) indicate that EXD2-deficient cells display poor ALT-EJ efficiency. (n= 3 independent experiments, statistical analysis was carried out using student's t-test, bars represent +/- SEM).

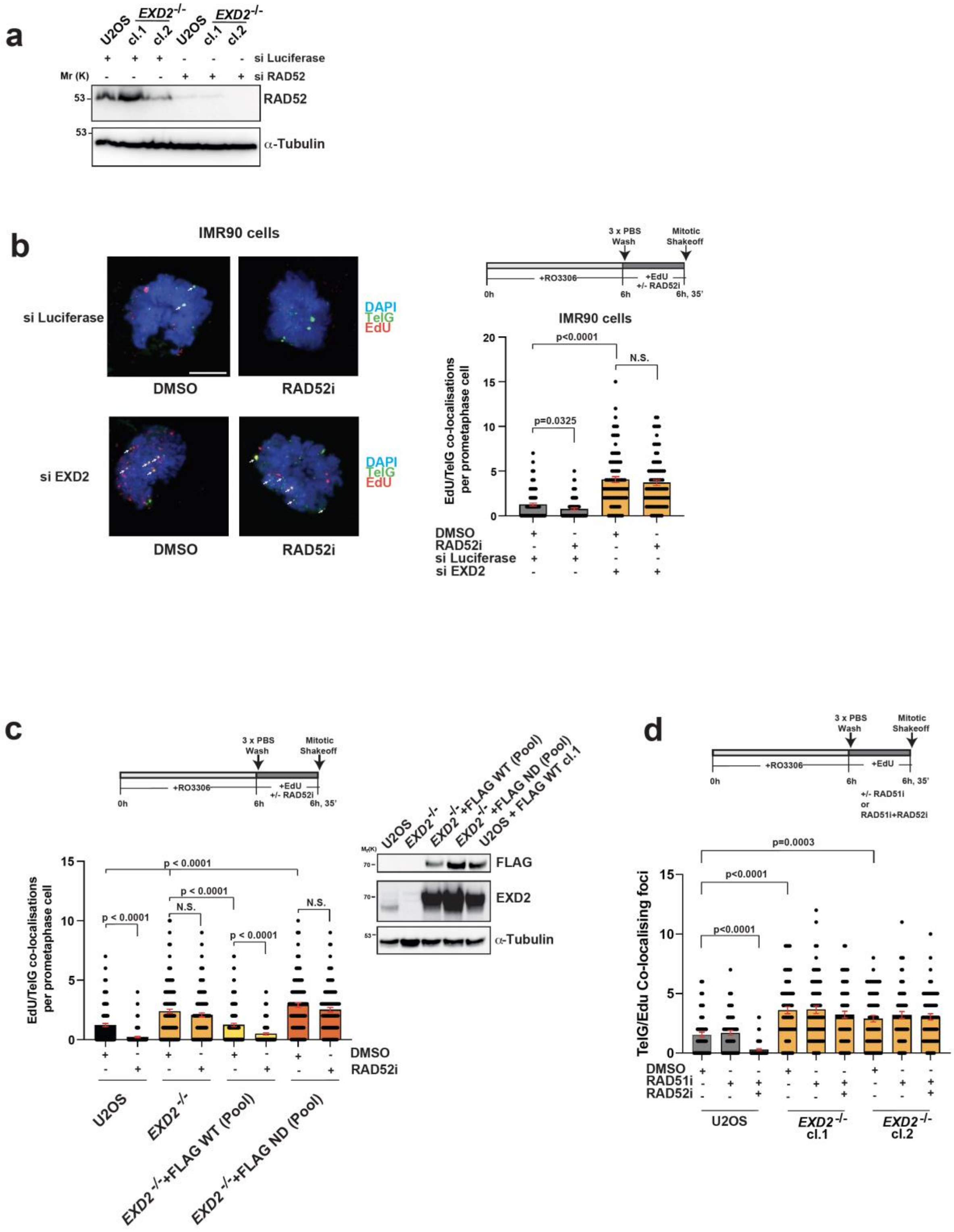


71 **Supplementary Figure 3**

**Supplementary Figure 3**

a) Confirmation of knockdown of RAD52 in WT U2OS and EXD2-deficient cells. $\alpha$-Tubulin acts as a loading control.

b) Immuno-FISH analysis in IMR90 cells treated with either control siRNA (si Luciferase) or siRNA targeting EXD2 as indicated. Cells were synchronised to G2 by RO-3306 followed by release to synchronised prometaphase in the presence of EdU as indicated. EdU incorporation was determined by Click-iT staining (red) and telomeres marked by a telomere-specific PNA FISH probe (TelG, green) with co-localisations indicated by arrows. DAPI acts as a DNA stain. Cells were treated with DMSO, or RAD52-inhibitor (AICAR), as indicated. (n= at least 84 prometaphases from 3 independent experiments, statistical significance was determined by Mann-Witney analysis, bars represent +/- SEM, scale bar = 10$\mu$m).

c) Immuno-FISH analysis in U2OS WT, *EXD2*$^{-/-}$ and pools of *EXD2*$^{-/-}$ cells stably expressing WT or nuclease dead (ND) FLAG-EXD2 as indicated. Cells were synchronised to G2 by RO-3306 followed by release to synchronised prometaphase in the presence of EdU as indicated. EdU incorporation was determined by Click-iT staining (red) and telomeres marked by a telomere-specific PNA FISH probe (TelG, green) with co-localisations indicated by arrows. DAPI acts as a DNA stain. Cells were treated with DMSO, or RAD52-inhibitor (AICAR), as indicated (n= at least 109 prometaphases from 3 independent experiments, statistical significance was determined by Mann-Witney analysis, bars represent +/- SEM, scale bar = 10$\mu$m). Western blotting confirms expression of WT and ND-FLAG EXD2. U2OS cells stably expressing WT FLAG EXD2 generated previously (U2OS +FLAG WT cl.1) act as a positive control for FLAG-EXD2 expression. $\alpha$-Tubulin acts as a loading control.

d) Immuno-FISH analysis in U2OS WT and *EXD2*$^{-/-}$ cells. Cells were synchronised to G2 by RO-3306 followed by release to synchronised prometaphase in the presence of EdU as indicated. EdU incorporation was determined by Click-iT staining (red) and telomeres marked by a telomere-specific PNA FISH probe (TelG, green) with co-localisations indicated by arrows. DAPI acts as a DNA stain. Cells were treated with either DMSO, a RAD51 inhibitor (B-02, 25μM) or RAD52 inhibitor (AICAR, 40μM), as indicated. (n= at least 64 prometaphases from 2 independent experiments, statistical significance was determined by Mann-Witney analysis, bars represent +/- SEM).

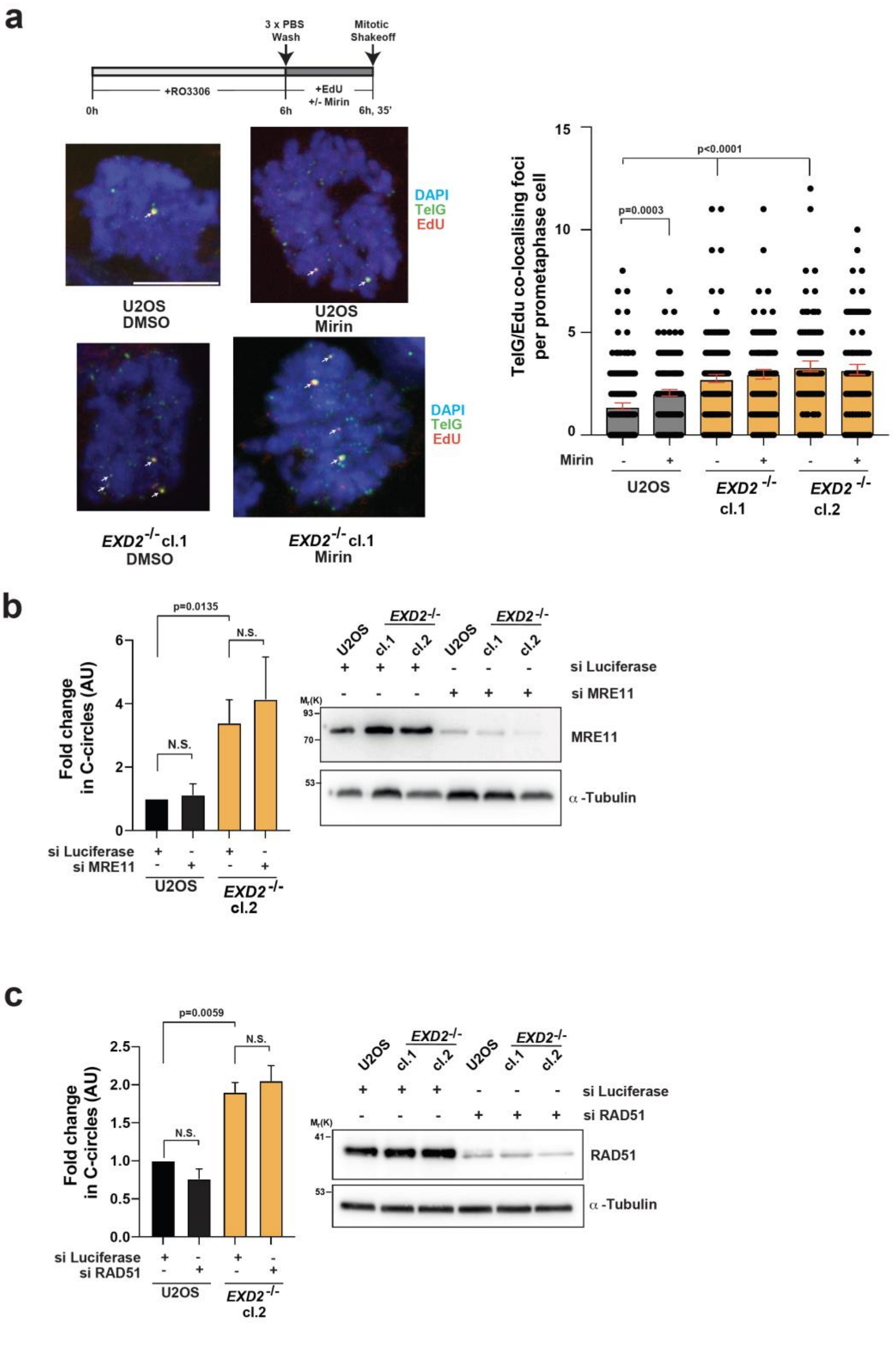


Supplementary Figure 4

**Supplementary Figure 4**

a) Analysis of EdU incorporation in isolated prometaphase cells from U2OS WT and EXD2-deficient clones treated with Mirin or mock treated with DMSO as indicated. EdU incorporation (Click-iT staining, red) colocalising with telomere-specific PNA FISH staining (TelG, green) is indicated with arrows. DAPI acts as a DNA stain (n= at least 80 prometaphases from 2 independent experiments, statistical significance was determined by Mann-Witney analysis, bars represent +/- SEM, scale bar = 10μm).

b) Analysis of C-circle abundance in U2OS and EXD2-deficient clones treated with control siRNA or siRNA targeting MRE11 as indicated (n= 8 measurements from 4 independent experiments, statistical significance was determined by student's t-test, bars represent +/-SEM). Western blotting confirming MRE11 knockdown upon siRNA treatment in WT U2OS and EXD2-deficient clones is included. α-Tubulin acts as a loading control.

c) Analysis of C-circle abundance in U2OS and EXD2-deficient clones treated with control siRNA or siRNA targeting RAD51 as indicated (n= 4 measurements from 2 independent experiments, statistical significance was determined by student's t-test, bars represent +/-SEM). Western blotting confirming RAD51 knockdown upon siRNA treatment in WT U2OS and EXD2-deficient clones is included. α-Tubulin acts as a loading control.

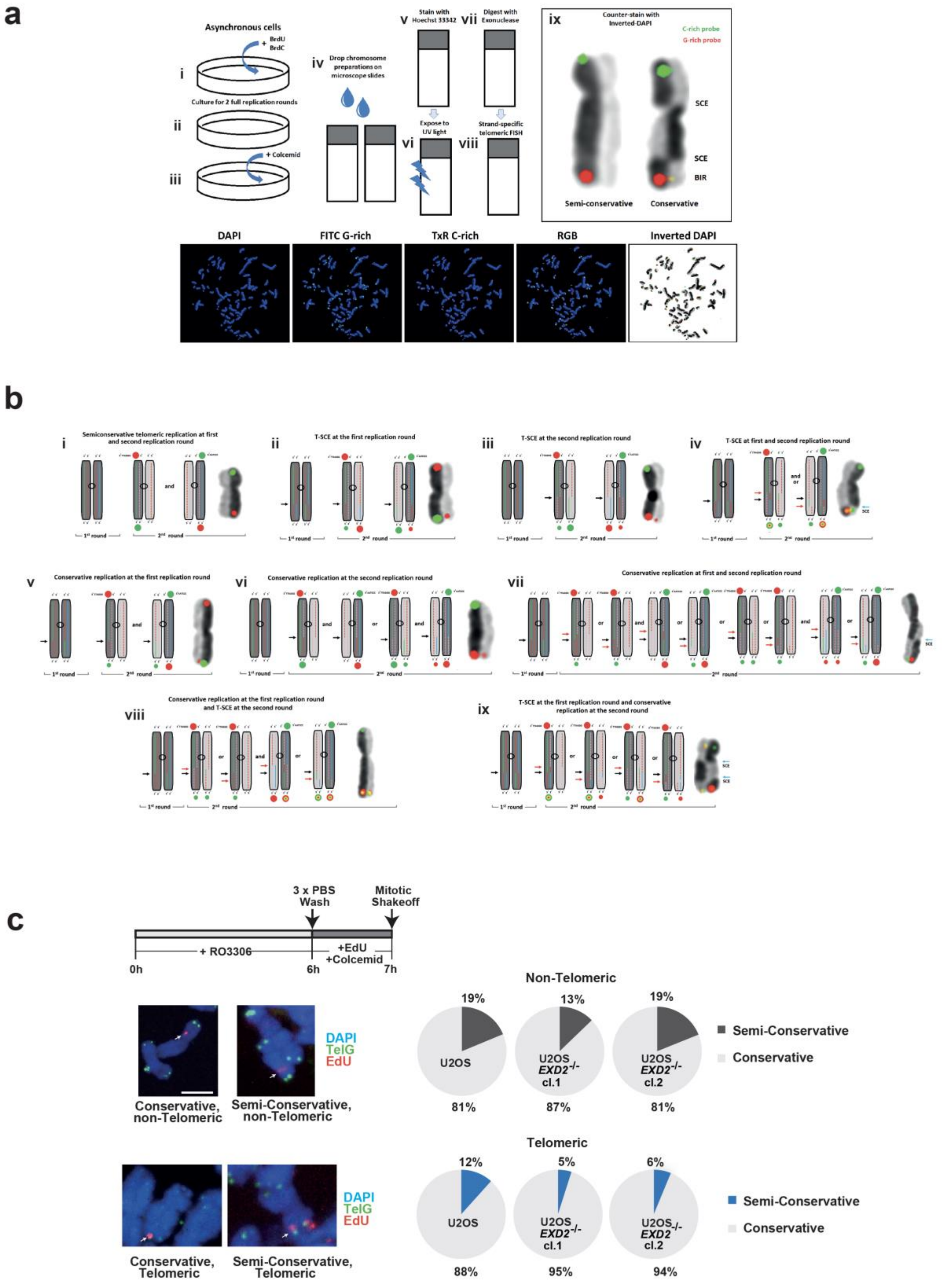


**Supplementary Figure 5**

**Supplementary Figure 5**

a) A modified protocol of segregated telomere strand-specific Chromosome Orientation Fluorescent In Situ Hybridization (CO-FISH), capable to quantify events of BIR-mediated conservative telomere replication per chromosome arm. The protocol entails living cells grown for two full replication rounds in presence of the synthetic Thymidine and Cytidine DNA base analogs, Bromodeoxyuridine (BrdU) and Bromodeoxycytidine (BrdC)(i-ii). Colcemid is added for 1-2 hours prior harvest to arrest dividing cells in mitosis (iii). Conventionally made chromosome preparations dropped onto microscope slides (iv), are then stained by Hoechst-33342 (v). Upon UV exposure, DNA crosslinking introduces nicks into the neosynthesized DNA strands throughout their full length and up to the telomeric termini (vi). Exonucleolytic digestion of nascent DNA (vii) is followed by non-denaturing telomere-strand-specific Peptide Nucleic Acid Analog (PNA) FISH (viii). A metaphase spread stained with segregated telomere strand-specific CO-FISH and inverted DAPI staining after two full replication rounds (ix). Differential labeling of sister chromatids allows detection of genomic Sister Chromatid Exchanges (SCE). Co-localized G-rich and C-rich telomeric signals at the tip of the light staining chromatid is always compatible with an event of BIR-mediated iconservative telomere synthesis.

b) Multiple possible scenarios of telomeric recombination in cultured ALT cells upon two replication rounds in presence of BrdU/BrdC. According to inverted DAPI staining, the darkly stained chromatids contain the original template DNA strands that did not incorporate BrdU/BrdC and are resistant to nucleolytic digestion (solid green lines indicate direction 3’-5’ and blue lines 5’-3’). Neosynthesized DNA strands that in the first replication round are depicted by red dotted lines and in the second round by pink dotted lines, will be degraded during the application of the CO-FISH protocol. Black arrows indicate primary breakpoints. Red arrows point to secondary breakpoints. Red and green spherical dots depict the expected staining pattern by telomere strand specific CO-FISH. Yellow dots indicate colocalization of G-rich and C-rich telomere fluorescence signals on the same chromatid tip. Due to highly increased telomeric length heterogeneity of the ALT cells, smaller red or green dots may range in fluorescence intensity from microscopically bright, to barely visible or invisible signals. Despite the variety of predicted phenotypes and the redundancy of several outcomes, co-localization of both template telomeric strands on the tips of the light staining chromatids by non-denaturing hybridization is always compatible with at least one recombinatorial event leading to conservative telomeric Break-Induced Replication occurring in the last two cell cycles (scenarios viii and ix). Matching segregated CO-FISH phenotypes are presented for each scenario. Note that some chromosomes underwent one or more genomic SCEs (blue arrows).

c) Analysis of EdU incorporation on isolated synchronised metaphase chromosomes harvested from U2OS WT and EXD2-deficient clones. EdU incorporation (Click-iT staining, red) in chromatid arms and at the telomere, marked with a telomere-specific FISH staining (TelG, green) are indicated. DAPI acts as a DNA stain. The percentage of observed semi-conservative (EdU incorporation on both sister chromatids) and conservative (EdU incorporation on one chromatid) is quantified for telomeric and non-telomeric events (n= at least 48 events from 3 independent experiments, scale bar = 5μm).

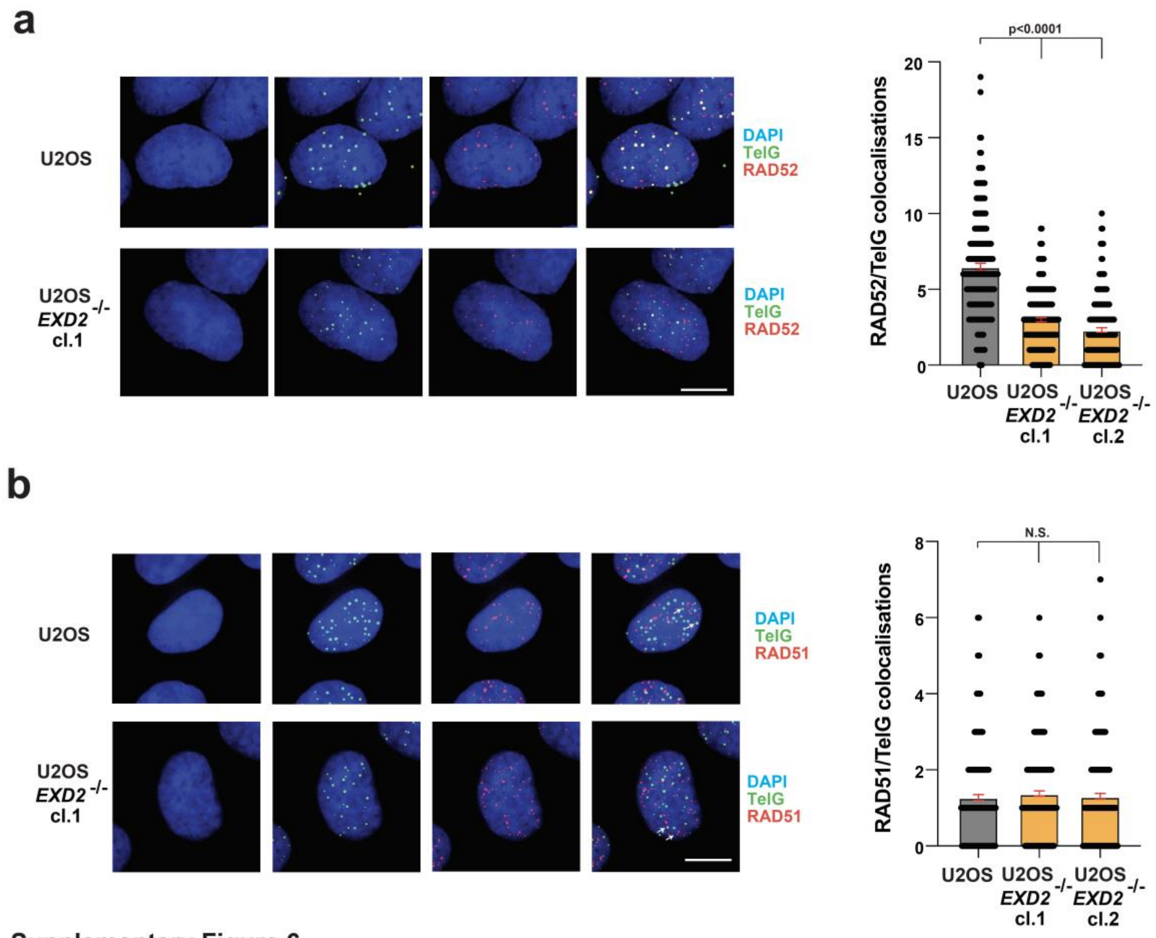


Supplementary Figure 6

**Supplementary Figure 6**

a) Immuno-FISH staining in WT and *EXD2*$^{-/-}$ U2OS cells using an antibody recognizing RAD52 (red) TelG PNA FISH probe staining (green) acts as a marker for the telomere and DAPI acts as a nuclear stain (n= at least 200 cells from 3 independent experiments, statistical analysis by Mann-Witney test, bars represent +/- SEM, scale bar = 10μm).

b) Immuno-FISH staining in WT and *EXD2*$^{-/-}$ U2OS cells using an antibody recognizing RAD51 (red) TelG PNA FISH probe staining (green) acts as a marker for the telomere and DAPI acts as a nuclear stain (n= at 300 cells from 3 independent experiments, statistical analysis by Mann-Witney test, bars represent +/- SEM, scale bar = 10μm).

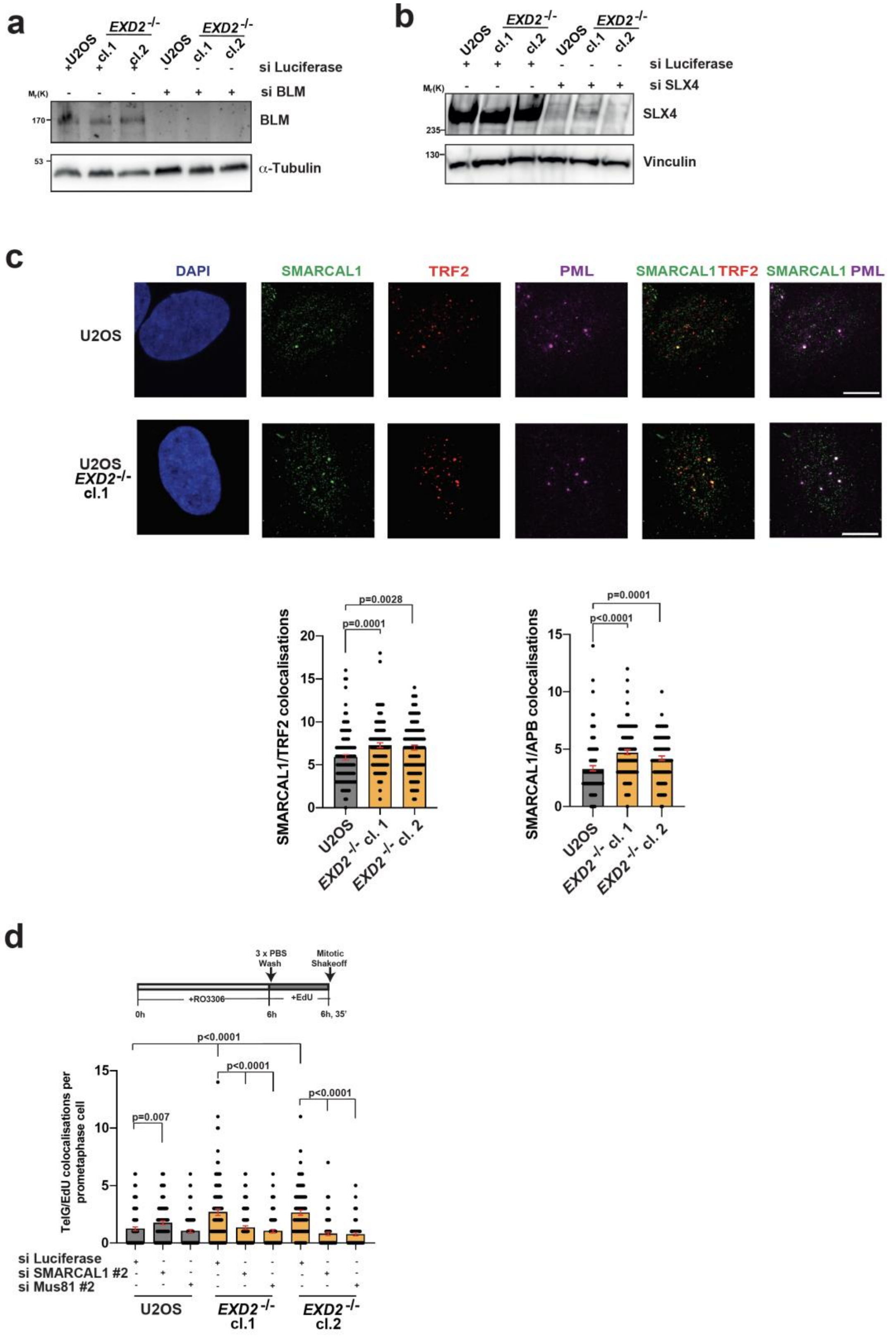


191 **Supplementary Figure 7**

**Supplementary Figure 7**

a) Western blotting confirming BLM knockdown in WT U2OS or *EXD2*$^{-/-}$ cells treated with either control siRNA or siRNA targeting BLM, as indicated. α-Tubulin acts as a loading control.

b) Western blotting confirming SLX4 knockdown in WT U2OS or *EXD2*$^{-/-}$ cells treated with either control siRNA or siRNA targeting SLX4, as indicated. Vinculin acts as a loading control.

c) Quantification of the localization of SMARCAL1 to telomeres and APBs by 4-colour immunofluorescence staining carried out in WT and *EXD2*$^{-/-}$ U2OS using antibodies raised against SMARCAL1 (green), TRF2 (red) and PML (magenta). DAPI acts as a nuclear stain (n= 100 cells from 2 independent experiments, statistical analysis by Mann-Witney test, scale bar = 10μm).

d) Analysis of EdU incorporation in isolated prometaphase cells from U2OS WT and EXD2-deficient clones treated with either control siRNA (si Luciferase) or siRNA targeting SMARCAL1 or MUS81, as indicated. EdU incorporation (Click-iT staining, red) co-localising with telomere-specific PNA FISH staining (TelG, green) is indicated with arrows. DAPI acts as a DNA stain (n= at least 67 prometaphases from 2 independent experiments, statistical significance was determined by Mann-Witney analysis, bars represent +/- SEM, scale bar = 10μm).

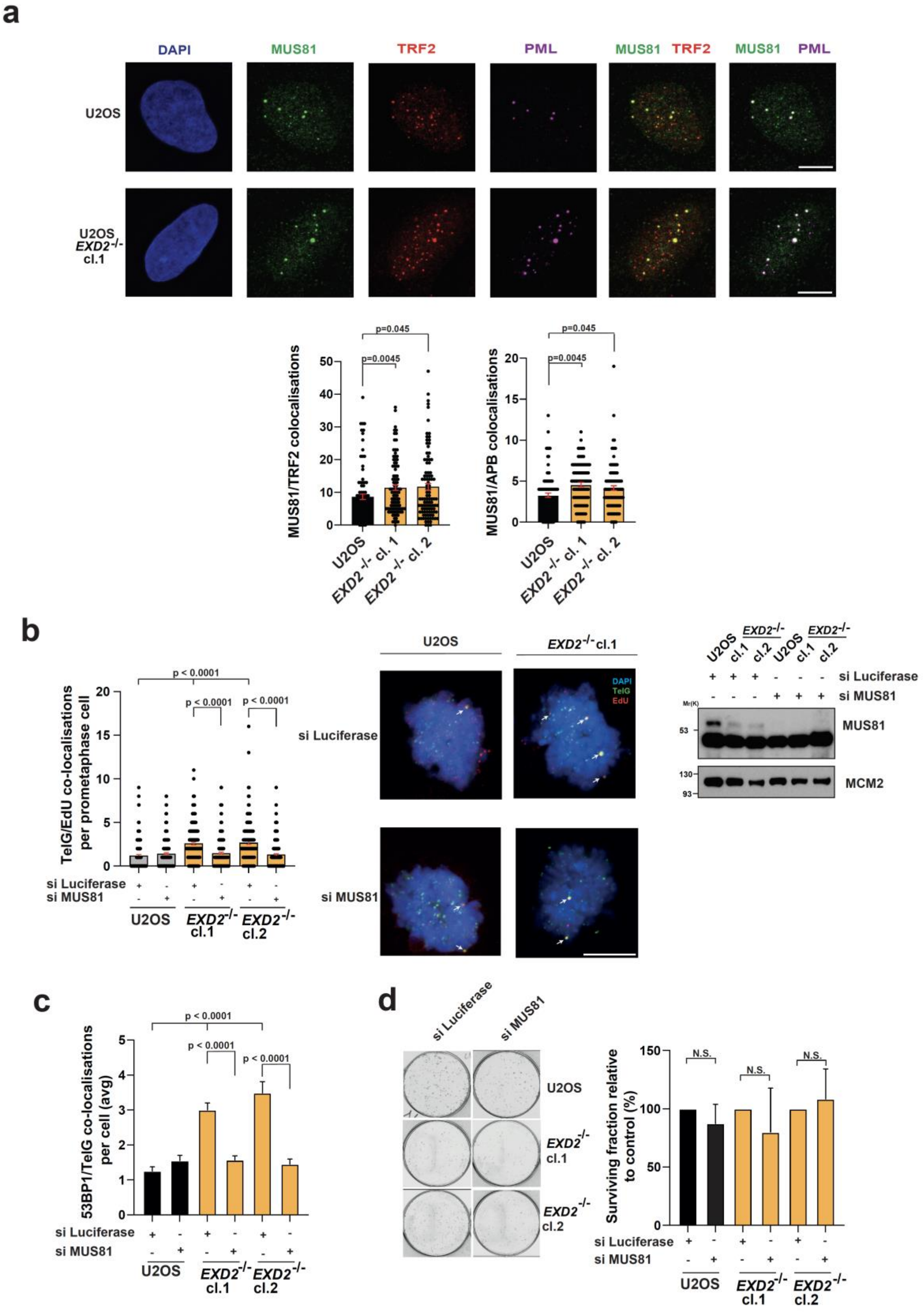


**Supplementary Figure 8**

**Supplementary Figure 8**

a) Quantification of the localization of MUS81 to telomeres and APBs by 4-colour immunofluorescence staining carried out in WT and *EXD2*$^{-/-}$ U2OS using antibodies raised against MUS81 (green), TRF2 (red) and PML (magenta). DAPI acts as a nuclear stain (n= 100 cells from 2 independent experiments, statistical analysis by Mann-Witney test, scale bar = 10μm).

b) Quantification of EdU co-localisations with TelG (G-rich telomere-specific PNA FISH probe in isolated synchronised prometaphase cells from U2OS and EXD2-deficient U2OS treated with control siRNA or siRNA targeting MUS81 (n= at least 150 prometaphases from 3 independent experiments, statistical significance was determined by Mann-Witney analysis, scale bar = 10μm. Confirmation of MUS81 knockdown by western blotting is presented. MCM2 acts as a loading control

c) Quantification of 53BP1 co-localisations with TelG (G-rich telomere specific probe) in WT or EXD2-deficient U2OS cells treated with control siRNA or siRNA targeting MUS81 (n= 150 cells from 3 independent experiments, statistical significance was determined by Mann-Witney analysis, bars represent +/- SEM).

d) Colony formation assay in U2OS WT or *EXD2*$^{-/-}$ cells treated with control siRNA or siRNA targeting MUS81, as indicated (n=3 independent experiments, bars represent +/-SEM).

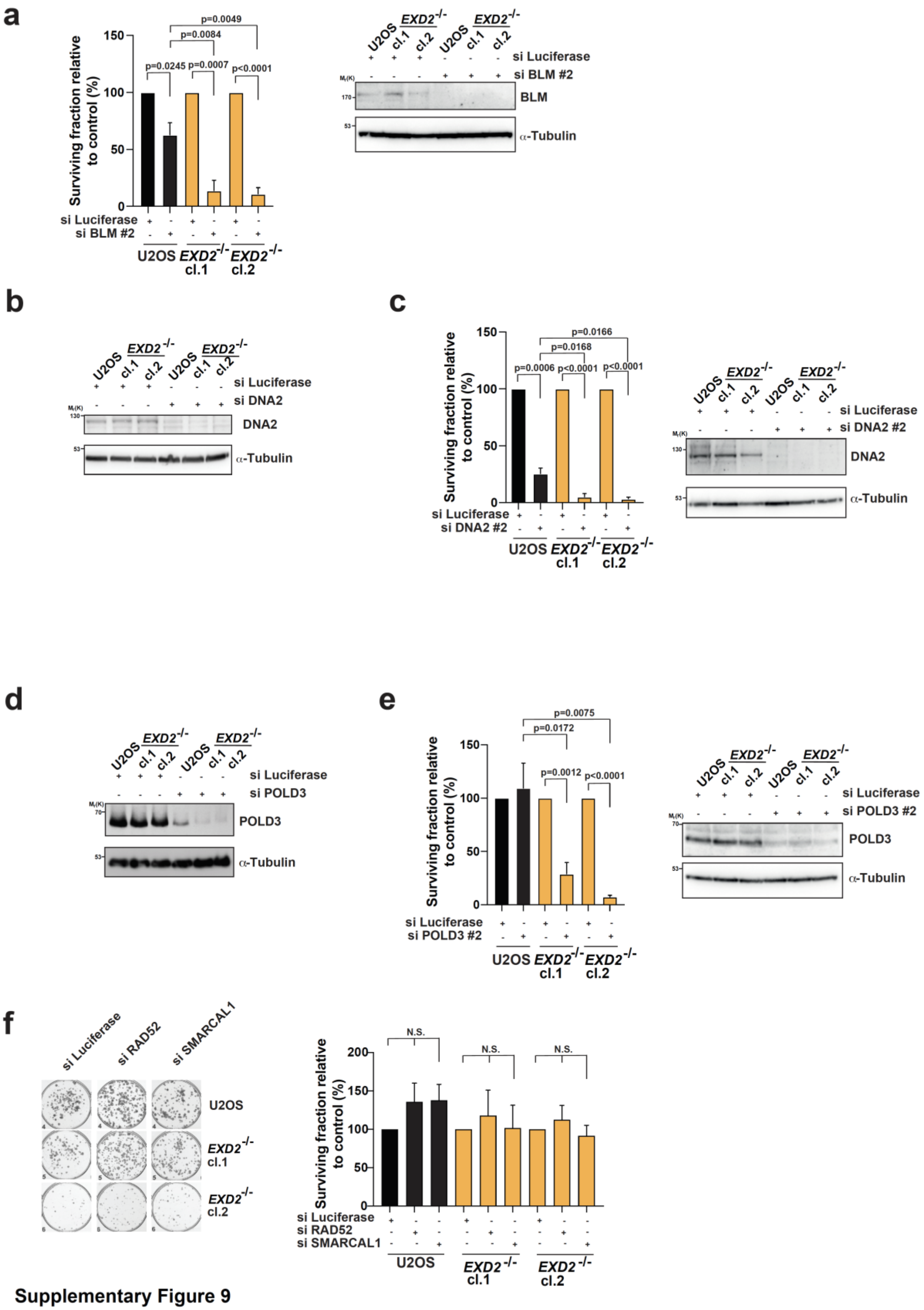


Supplementary Figure 9

**Supplementary Figure 9**

a) Colony formation assays in WT U2OS and *EXD2*$^{-/-}$ cells treated with siRNA targeting BLM or control siRNA (Luciferase). Surviving fraction relative to control is quantified (n= 5 independent experiments, statistical significance was determined by student's t-test, bars represent +/-SEM). Confirmation of BLM knockdown by western blotting is presented. $\alpha$-Tubulin acts as a loading control.

b) Confirmation of DNA2 knockdown in U2OS WT and *EXD2*$^{-/-}$ cells treated with siRNA targeting DNA2 or control siRNA (Luciferase), as indicated. $\alpha$-Tubulin acts as a loading control.

c) Colony formation assays in WT U2OS and *EXD2*$^{-/-}$ cells treated with siRNA targeting DNA2 or control siRNA (Luciferase). Surviving fraction relative to control is quantified. (n= 4 independent experiments, statistical significance was determined by student's t-test, bars represent +/-SEM). Confirmation of DNA2 knockdown by western blotting is presented. $\alpha$-Tubulin acts as a loading control.

d) Confirmation of POLD3 knockdown in U2OS WT and *EXD2*$^{-/-}$ cells treated with siRNA targeting POLD3 or control siRNA (Luciferase), as indicated. $\alpha$-Tubulin acts as a loading control.

e) Colony formation assays in WT U2OS and *EXD2*$^{-/-}$ cells treated with siRNA targeting POLD3 or control siRNA (Luciferase). Surviving fraction relative to control is quantified. (n= 6 technical repeats from 3 independent experiments, statistical significance was determined by student's t-test, bars represent +/-SEM). Confirmation of POLD3 knockdown by western blotting is presented. $\alpha$-Tubulin acts as a loading control.

f) Colony formation assay in U2OS WT or *EXD2*$^{-/-}$ cells treated with control siRNA or siRNA targeting RAD52 or SMARCAL1, as indicated (n=3 independent experiments, bars represent +/-SEM).

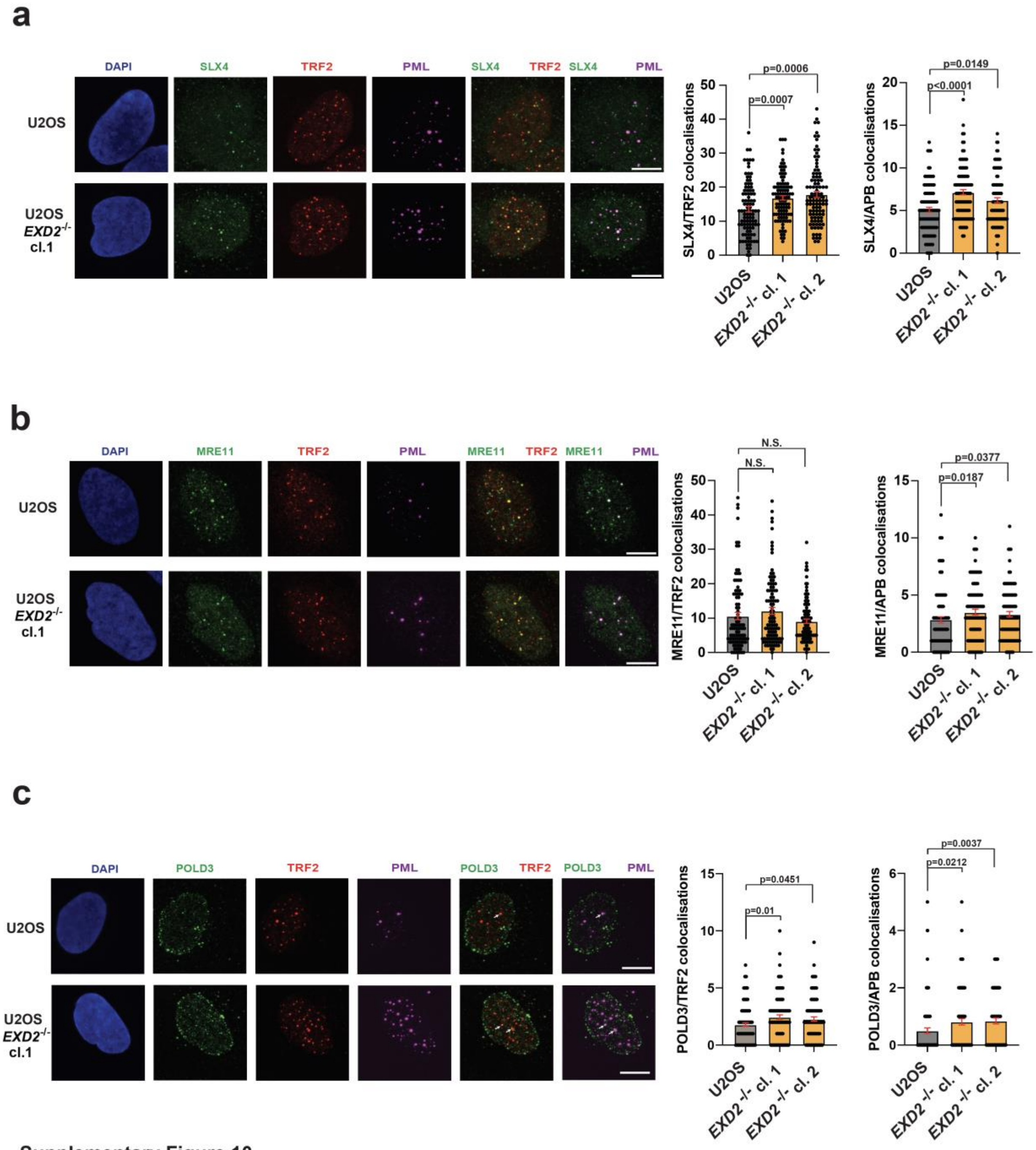


**Supplementary Figure 10**

**Supplementary Figure 10**

a) Quantification of the localization of SLX4 to telomeres and APBs by 4-colour immunofluorescence staining carried out in WT and *EXD2*$^{-/-}$ U2OS using antibodies raised against SLX4 (green), TRF2 (red) and PML (magenta). DAPI acts as a nuclear stain (n= 100 cells from 2 independent experiments, statistical analysis by Mann-Witney test, scale bar = 10$\mu$m).

b) Quantification of the localization of MRE11 to telomeres and APBs by 4-colour immunofluorescence staining carried out in WT and *EXD2*$^{-/-}$ U2OS using antibodies raised against MRE11 (green), TRF2 (red) and PML (magenta). DAPI acts as a nuclear stain (n= 100 cells from 2 independent experiments, statistical analysis by Mann-Witney test, scale bar = 10$\mu$m).

c) Quantification of the localization of POLD3 to telomeres and APBs by 4-colour immunofluorescence staining carried out in WT and *EXD2*$^{-/-}$ U2OS using antibodies raised against POLD3 (green), TRF2 (red) and PML (magenta). DAPI acts as a nuclear stain (n= 100 cells from 2 independent experiments, statistical analysis by Mann-Witney test, scale bar = 10$\mu$m).
